# Senolytic Therapy as a Preventive Strategy for Low Back Pain

**DOI:** 10.1101/2025.10.08.681005

**Authors:** Saber Ghazizadeh, Hosni Cherif, Matthew Mannarino, Juiena Sagir, Magali Millecamps, Jean A. Ouellet, Laura S. Stone, Lisbet Haglund

## Abstract

Cell senescence drives inflammation and tissue breakdown and is a key hallmark of aging. Low back pain is strongly linked to age-related degeneration of spine tissues, and with an accumulation of senescent. Here we show that preventive administration of the senolytic agents o-vanillin and RG-7112 prevent the development of pain-related behaviour in young *sparc^-/-^* mice. Treated mice exhibit a reduction of senescence markers in the intervertebral discs, vertebral endplates, vertebral bone, and spinal cord, alongside a dampening of pro-inflammatory senescence-associated secretory factors in these tissues. This early senolytic intervention also preserves intervertebral disc volume and vertebral bone microarchitecture, indicating protection against structural degeneration of the spine. These findings demonstrate that targeting cellular senescence at an early stage can mitigate degenerative changes and pain, supporting senolytic therapy as a promising preventive strategy for musculoskeletal decline.

## INTRODUCTION

Cellular senescence, a key hallmark of aging and degenerating tissues, is characterized by a stable and essentially irreversible cell-cycle arrest that develops in response to stressors such as DNA damage, oxidative stress, mitochondrial dysfunction, and epigenetic instability (1, 2). Senescent cells (SnCs) accumulate in tissues over time and actively contribute to age-related pathologies including osteoarthritis (3–5), neurodegeneration (6–8), and cancer (9–11). Beyond loss of proliferative potential, SnCs develop a pro-inflammatory phenotype referred to as the senescence-associated secretory phenotype (SASP) (9, 12). The SASP is composed of cytokines, chemokines, proteases, growth factors, and matrix-degrading enzymes (13, 14). These molecules alter the surrounding tissue environment by impairing structural integrity, amplifying inflammation, and promoting paracrine senescence, thereby exacerbating tissue dysfunction. Due to their resistance to apoptosis, SnCs persist in tissues and act as active drivers of tissue deterioration (15, 16).

Identification of senescent cells has relied on SA-β-gal lysosomal activity (17); however, this marker lacks specificity, as elevated β-gal activity can occur in non-senescent contexts, including certain immune, quiescent or stressed cells (18). More definitive indicators include the cyclin-dependent kinase inhibitors *p21* and *p16^Ink4a^*, which are broadly accepted as key molecular markers of early and late senescence, respectively (19–21). While *p21* is primarily induced by p53 and reflects transient or early senescence in response to acute stress (22), *p16^Ink4a^*tends to accumulate with age and marking a more stable, late-stage senescent phenotype (20, 23). These subtypes also differ in their SASP composition and apoptotic threshold, with cells expressing high *p16^Ink4a^* levels being more resistant to clearance (24). Understanding this heterogeneity is crucial for designing senolytic strategies, as certain drugs preferentially target either *p16^Ink4a^* or *p21*-expressing senescent cells, influencing treatment outcomes in diseases.

Pharmacological removal of senescence burden with senomorphics and senolytic agents has emerged as a promising therapeutic approach for delaying or reversing tissue deterioration (25, 26). Drugs such as navitoclax (a Bcl-2 inhibitor), FOXO4-DRI, and the MDM2 inhibitor RG-7112 exemplify distinct senolytic classes. Natural compounds, including quercetin, fisetin, and o-vanillin, also exhibit senolytic or senomorphic activity in various models (3, 10, 25–32). Senomorphics are drugs designed to suppress the detrimental effects of SASP components without killing SnCs (32, 33). Senolytic drugs selectively target and kill SnCs. Class I of senolytics target the apoptosis-primed BH3 networks of senescent cells; Class II of senolytics inhibit survival pathways elicited by senescent cells, and class III that disrupt homeostatic processes already challenged in senescent cells (34).

Low back pain (LBP), often associated with intervertebral disc (IVD) degeneration, is the leading cause of years lived with disability globally (35–38). The socioeconomic impact is substantial, with healthcare costs in the United States alone surpassing $100 billion annually (39, 40). Degenerating discs display key features of cellular senescence, including elevated expression of *p16^Ink4a^*, *p21*, and SASP components such as IL-1β, IL-6, and matrix metalloproteinases (41–45). These molecules contribute to extracellular matrix breakdown, neurovascular infiltration (46), and nociceptor sensitization (47), ultimately driving both structural failure and chronic pain (48).

In our previous study, we demonstrated that treatment with either o-vanillin or RG-7112 eliminated SnCs and reduced SASP factors in human IVD cells and tissues. These treatments improved matrix homeostasis and cell viability, indicating translational potential (49–54).

We have also previously demonstrated that o-vanillin and RG-7112 significantly reduced the SnC burden, improved disc structure, suppressed inflammatory mediators, and attenuated behavioural signs of back pain in middle-aged *sparc^-/-^*mice, which had well-established IVD degeneration and back pain (54).

RG-7112 is a synthetic MDM2 inhibitor that functions by disrupting the MDM2–p53 interaction, sending SnCs to apoptosis (55). The natural flavonoid o-vanillin is a curcumin-derived metabolite that possesses both senolytic and senomorphic properties (49, 53, 56). The senomorphic activity is mediated partially through modulation of Nrf2 and NF-κB signalling (49).

Building on our prior findings, this study investigates whether administration of o-vanillin and RG-7112 can prevent or delay IVD degeneration and associated pain in young *sparc^-/-^* mice at the time when signs of IVD degeneration and back pain are emerging. In addition, we assessed whether combination therapy would provide enhanced benefit over single therapy, given the distinct molecular mechanisms of these agents. Our study was designed to evaluate the impact of preventative senotherapeutic administration on disc and bone health, inflammatory signalling, and pain behaviour.

## Results

### Senotherapeutics prevented the development of a pain phenotype in *sparc^-/-^* mice

*Sparc^-/-^* mice begin to show signs of LBP at 4 months of age, with pain becoming well-established by 7 months (57–59). To follow the onset and progression of LBP phenotype in the *sparc^-/-^* mice, we conducted a comprehensive behavioural analysis once a month between 4 and 9 months of age, in both male and female mice. Experimental groups and treatment timelines are depicted in Figure 1A. *Sparc^-/-^* mice received weekly oral treatments with vehicle, o-vanillin (O, 100 mg/kg), RG-7112 (R, 5 mg/kg), or a combination (O+R, 100 mg/kg + 5 mg/kg). Treatment dosages were based on earlier experiments conducted in middle-aged *sparc^-/-^* mice (54).

Grip strength was used to determine axial pain, while von Frey and acetone-evoked behaviour were used to evaluate radiating pain. Vehicle-treated *sparc^-/-^* mice developed a progressively worsening pain phenotype across all tests, whereas wild-type (wt) mice maintained a stable, non-painful behavioural profile throughout the experimental period.

At baseline, no significant differences were detected between wt and *sparc^-/-^* mice in cold hypersensitivity measured by the acetone-evoked behaviour tests (Figure 1B-D). However, a small but significant difference was detected at baseline between the groups in grip strength (Figure 1E-G) and mechanical sensitivity (Figure 1H-J).

Treatment with o-vanillin or RG-7112 significantly attenuated the development of cold hypersensitivity, indicative of radiating pain. The effect was notable after 2 months of o-vanillin and 1 month of RG-7112 treatment. After 4 months of o-vanillin and 5 months of RG-7112 treatment, cold sensitivity was comparable to that observed in *wt* animals (Figure 1B–D). In the grip strength test, indicative of axial discomfort, untreated *sparc^-/-^* mice exhibited a reduction in grip strength over time. This decline was prevented after 3 months of o-vanillin and 2 months of RG-7112 treatment; however, values remained below the wt levels (Figure 1E–G). Mechanical sensitivity was assessed by the von Frey threshold test. The decline in mechanical sensitivity, reflecting increased radiating pain, was significantly reduced after four months of treatment with either o-vanillin or RG-7112, with both groups reaching *wt*-equivalent thresholds after 4 months of treatment (Figure 1H–J). By the fifth month, both senotherapeutic agents elicited significantly less pain behaviour across the three tests (Figure 1B-C, E-F, H-I).

To investigate the potential of a synergistic or additive effect, the drugs were administered as a combination treatment. After 5 months of treatment, combination therapy significantly improved grip strength (Figure 1E-F) and mechanical sensitivity (Figure 1H-I), compared to the drug administered as a single treatment. Cold sensitivity, on the other hand, was not further improved by a combination treatment (Figure 1B-D). There were no significant differences between male and female animals. These findings demonstrate that individual treatment with either o-vanillin or RG-7112 effectively prevents the development of low back pain, while a combination treatment further improves grip strength and reduces mechanical sensitivity.

Male and female animals are shown together as no significant sex differences were found. We found no difference in the body weight and mortality between the groups (Figure S1A, B and C).

**Figure 1.**
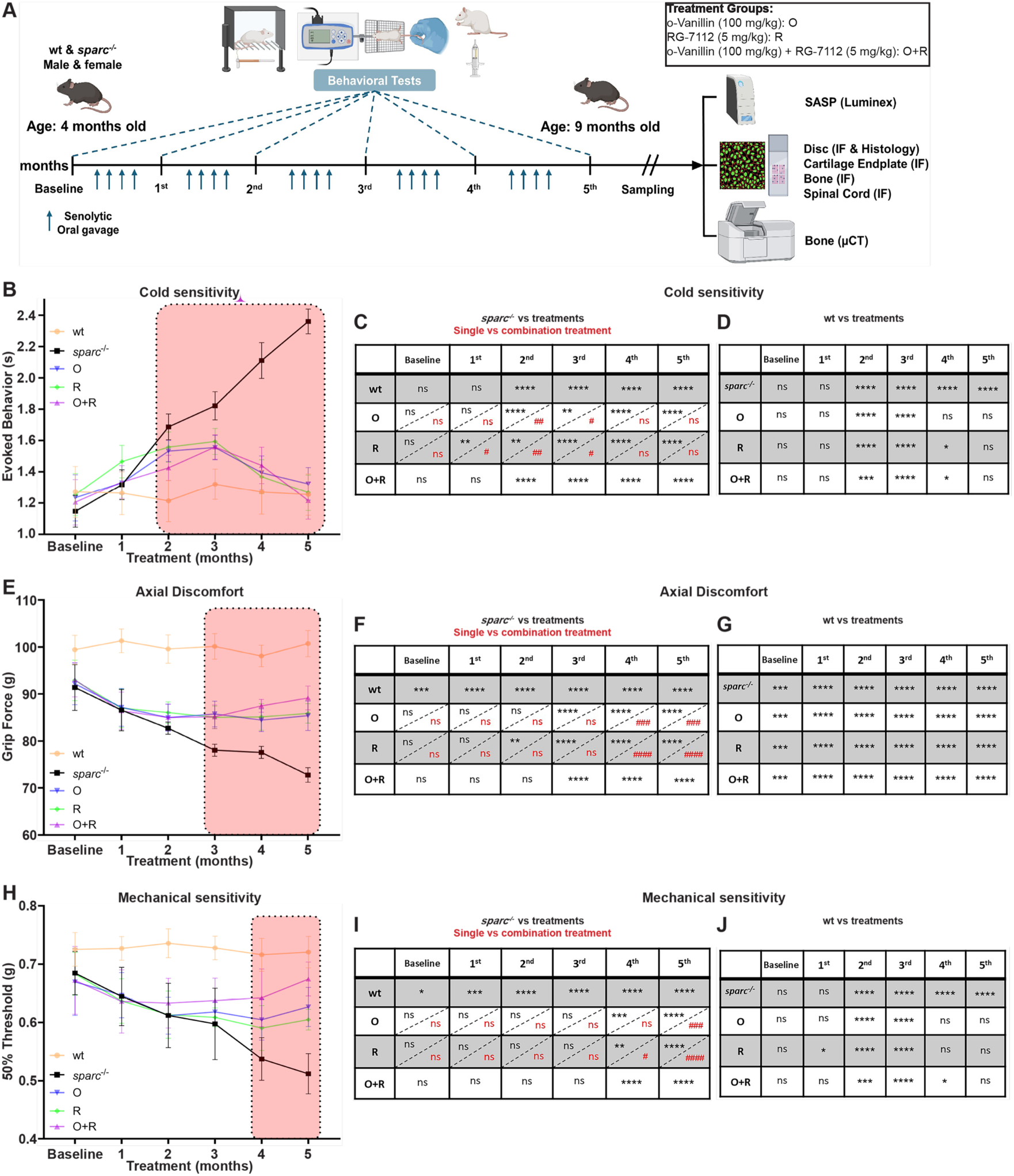
Preventive senolytic therapy attenuated pain-related behaviour in *sparc*^-/-^ mice. (**A**) Experimental design and treatment groups. *Sparc^-/-^* mice received weekly oral gavage of vehicle or senolytic treatment: o-vanillin (O), RG-7112 (R) or combination (O+R). Pain assessment tests were performed every month (grip strength, acetone-evoked behaviour, and von Frey); μCT, micro–computed tomography; IF, immunofluorescence; SASP, senescence-associated secretory phenotype. (**B-D**) Cold sensitivity measurements and comparisons (C) between *sparc*^-/-^ and treatments, (D) between *wt* and treatments; (**E-G**) Grip strength measurements and comparisons (F) between *sparc*^-/-^ and treatments, (G) between *wt* and treatments; (**H-J**) Mechanical sensitivity measurements and comparisons (I) between *sparc*^-/-^ and treatments, (J) between *wt* and treatments. *n* = 15 animals per group (7 males and 8 females). Data are represented as means ± SEM and analyzed by one-way ANOVA with Tukey’s post hoc test. */#*P* < 0.05, **/##*P* < 0.01, ***/###*P* < 0.001, ****/#### *P* < 0.0001 and ns No significant. * Indicate a significant difference compared with *sparc*^-/-^ or wt, and # indicates a significant difference comparing single with combination treatment. Schematic in (A) was generated using Bio Render.

### Senotherapeutics decreased SnCs and pain mediators in the spinal cord

The presence of SnCs in the central nervous system (CNS), including neural and glial populations, has been implicated in the development of chronic pain (60, 61). We found a significantly higher number of *p16^Ink4a^*-positive cells in the dorsal horn of vehicle-treated *sparc*^-/-^ mice compared to the *wt* group, consistent with previous reports (Figure 2A-B). Treatment with o-vanillin, RG-7112 or their combination all resulted in significantly fewer *p16^Ink4a^*-positive cells compared to the vehicle-treated *sparc*^-/-^ group, and in all cases, levels were comparable to those observed in *wt* controls (Figure 2B).To further assess the role of senolytics in modulating pain phenotype and to identify which cell populations were senescent, we colocalized the expression of *p16^Ink4a^* with the glial fibrillary acidic protein (GFAP), a marker of reactive astrocytes (Figure 2C), ionized calcium-binding adaptor molecule 1 (Iba1), a marker of activated microglia (Figure 2E), and neuronal nuclei (NeuN) marker (Figure 2G). S*parc*^-/-^ mice displayed a higher number of senescent GFAP-positive astrocytes (Figure 2D, I), Iba1-positive microglia (Figure 2F, J) and NeuN-positive neurons (Figure 2H, K) per area compared to the wt group.

Senolytic treatment with either o-vanillin or RG-7112 significantly reduced the number of senescent GFAP-positive astrocytes, Iba1-positive microglia, and senescent NeuN-positive neurons in the dorsal horn (Figure 2I-K). The combination did not provide an added effect compared to single drug treatment. Interestingly, all treatments restored levels to those observed in wt controls. The total number of *p16^Ink4a^* cells exceeded the number of cells identified through colocalization, indicating that additional cell types may contribute to the overall senescence burden.

**Figure 2.**
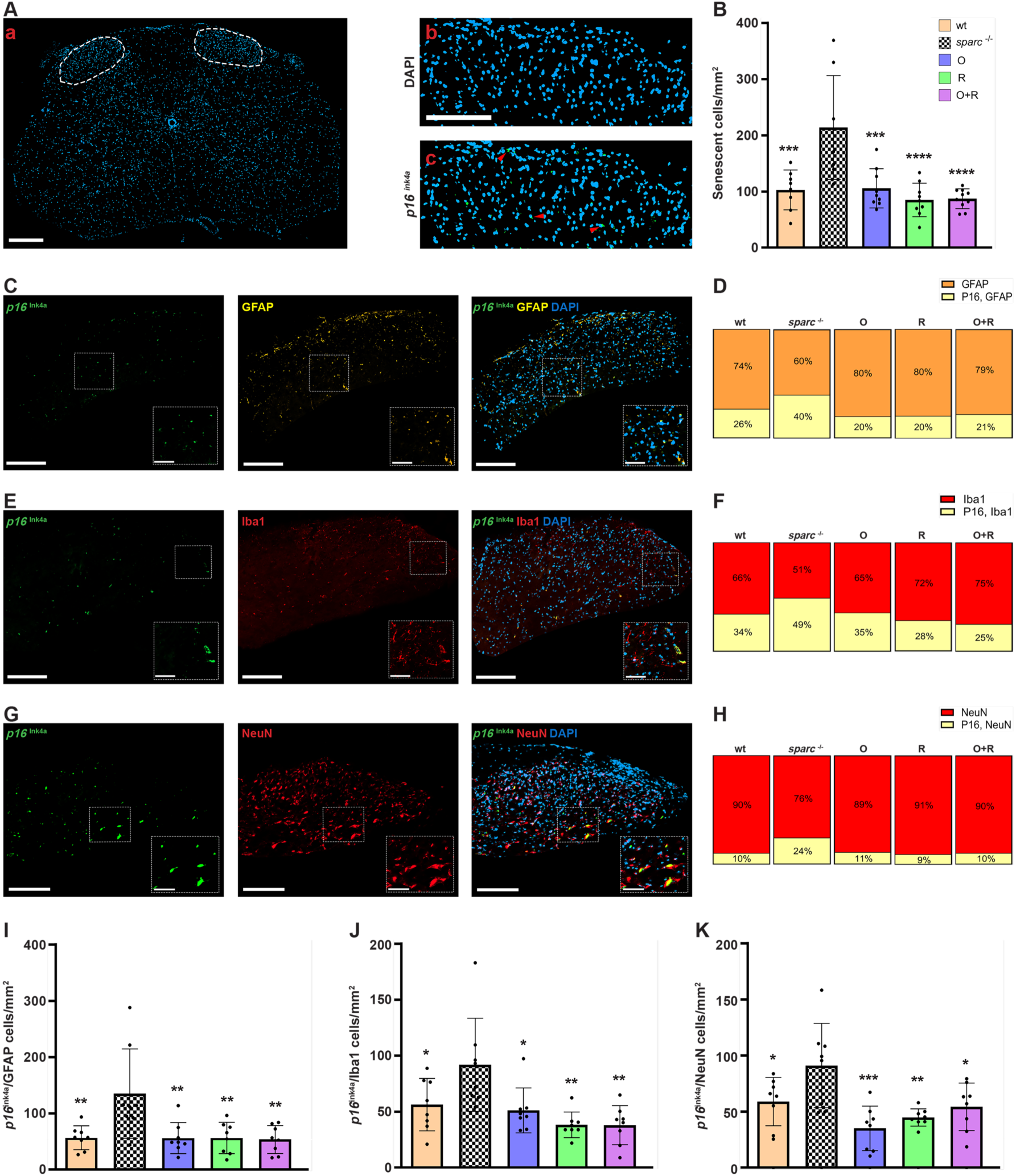
Senolytic treatment reduced senescent cell in the dorsal horn. (**A**) (a) Representative image of a DAPI-stained spinal cord section with the dorsal horn outlined with a dashed line and (b-c) DAPI and *p16^Ink4a^* immunoreactivity in the dorsal horn. (**B**) Quantification of *p16^Ink4a^*-positive cells in the dorsal horn. (**C**) Representative image showing*p16^Ink4a^*, and GFAP expression. **(D)** Percentage of GFAP and GFAP/ *p16^Ink4a^*double-labelled cells. **(E)** Representative image showing *p16^Ink4a^*and Iba1 expression. **(F)** Percentage of Iba1 and Iba1/ *p16^Ink4a^* double-labelled cells. **(G)** Representative image showing *p16^Ink4a^*, and NeuN expression. **(H)** Percentage of NeuN and NeuN/ *p16^Ink4a^* double-labelled cells. Number of cells/ mm^2^. N= 8-10 animals per group (4 to 5 males or females). DAPI served as a nuclear counterstain. For each animal, the mean of three independent images was calculated for group analysis. Scale bars = 200 μm and 80 μm. Data are presented as means ± SD and were analyzed by an ordinary one-way ANOVA followed by Tukey’s post hoc test. \**P* < 0.05, \*\**P* < 0.01, \*\*\**P* < 0.001, and \*\*\*\**P* < 0.0001.* Indicates a significant difference compared with *sparc^-/^*^-^.

### Senotherapeutics reduced SASP factor release from the IVDs

IVDs from *sparc^-/-^* mice exhibit elevated secretion of SASP factors, which have been previously associated with LBP and disc degeneration (54). In our prior study, we demonstrated that by 7 and 9 months, *sparc^-/-^* IVDs display increased expression of multiple SASP components, and that therapeutic intervention with o-vanillin and RG-7112 attenuated SASP release (54). However, it remains unknown whether early intervention can prevent the establishment of a senescence-associated inflammatory microenvironment.

To investigate the effect of preventative treatment on SASP factor release, lumbar IVDs from treated and untreated animals were harvested at the termination of the experiment. The lumbar region was selected due to its critical biomechanical role and its high susceptibility to age-related degeneration and LBP in both humans and *sparc^-/-^* mice (58). The secretion profiles of 15 SASP-associated cytokines, chemokines and growth factors were quantified using a Luminex multiplex assay (Figure 3A–O).

As anticipated, vehicle-treated *sparc^-/-^* IVDs exhibited significantly elevated levels of all 15 SASP factors compared to *wt* control. Treatment with either o-vanillin, RG-7112 or combination led to significant reductions in all 15 SASP factors relative to vehicle-treated *sparc^-/-^*mice. Notably, combination treatment yielded a statistically additive effect only for IL-6, with no additional benefit observed for the remaining factors when compared to single treatments. Among the 15 SASP factors measured, levels of eight (CXCL1, IL-1β, IL-2, CXCL10, M-CSF, CXCL9, RANKL, and VEGF-α) were not statistically different from those observed in *wt* controls. A summary of the measured concentrations and statistical comparisons is provided in Table S2.

**Figure 3.**
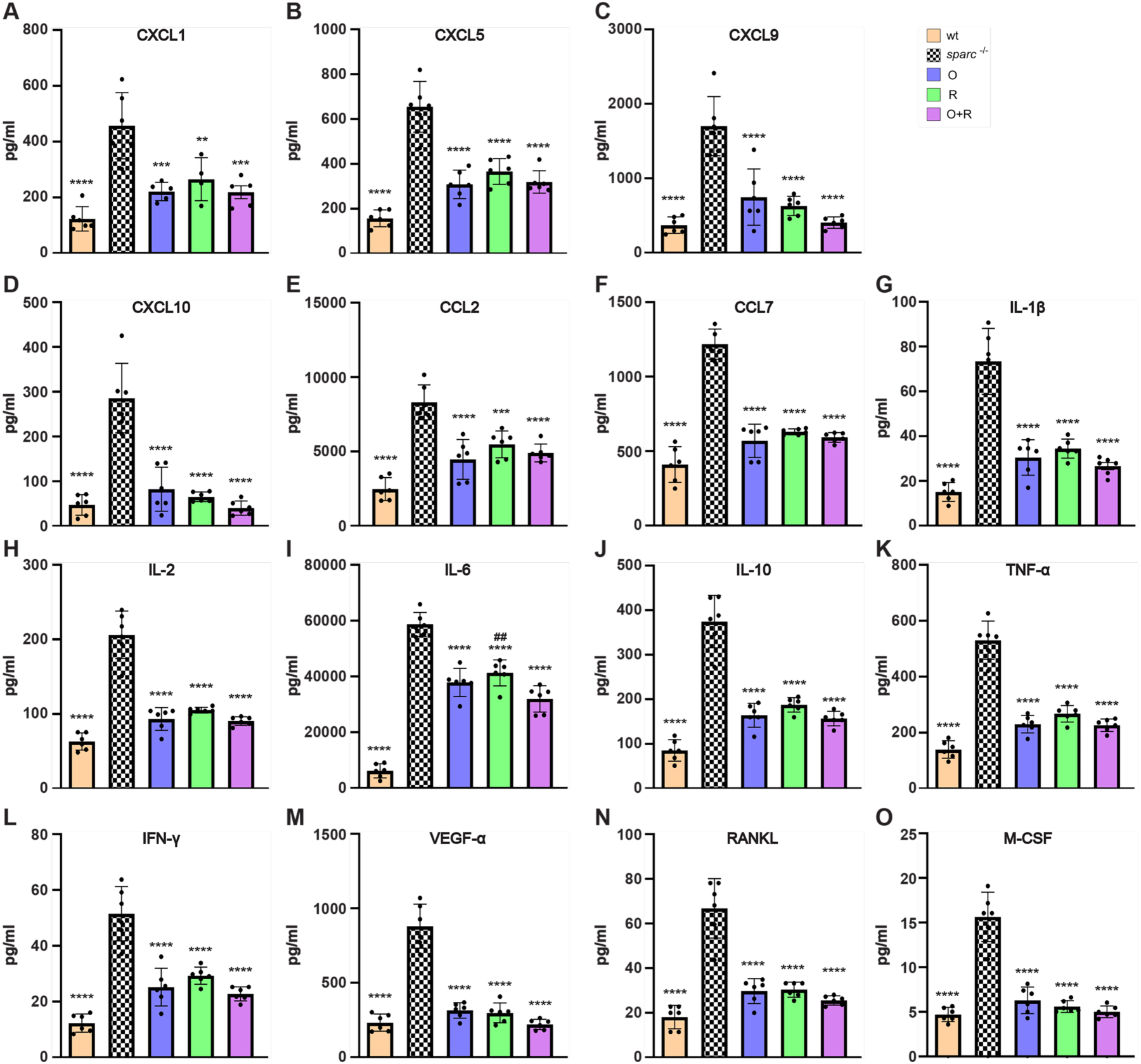
Senotherapeutic treatment decreased SASP factor secretion from the IVDs. (**A** - **F**) chemokines, (**G**-**L**) cytokines, and (**M**-**O**) growth factor release from IVDs was assessed by a multiplex assay. The data is presented as means ± SD; one-way ANOVA and post hoc comparison Tukey’s were used to measure significant differences between the groups. ##*P* < 0.01, \*\*\**P* < 0.001, and \*\*\*\**P* < 0.0001. * Indicates a significant difference compared to *sparc*^-/-^, and # indicates a significant difference compared to combination treatment. Four discs represent one measure per animal. *n* = 6 animals (3 males and 3 females) in each group.

### Senotherapeutics prevented the accumulation of SnCs, and loss of disc volume and resulted in a reduced disc degeneration score

Next, we evaluated whether preventative senolytic treatment could preserve disc volume and IVD health. Previous results where animals with well-established degeneration were treated showed that the combination treatment was significantly better than single drugs (54). Here we confirmed a significantly lower IVD volume in *sparc^-/-^* compared to *wt* mice (Figure 4A-B). Treatment with the drugs individually resulted in a slightly preserved disc volume; however, these changes reached statistical significance only following RG-7112 treatment. Importantly, the combination therapy improved the outcome and preserved disc volume to a higher and more significant level, preserving IVD volume to values comparable to those of wt mice (Figure 4A, B).

Then, the extent of IVD degeneration was assessed through histological grading using a validated scoring system (54, 57). Treatment with either o-vanillin or RG-7112 resulted in a significantly improved IVD degeneration score compared to vehicle-treated *sparc*^-/-^. Notably, combination therapy did not yield an added effect. Furthermore, despite these improvements, none of the treatment groups achieved degeneration scores comparable to those of *wt* controls (Figure 4C-D).

In addition to assessing IVD degeneration scores, we evaluated the effect of senolytic treatment on the accumulation of SnCs by quantifying the number of *p16^Ink4a^*and *p21*-positive IVD cells. Our previous studies have demonstrated that systemic administration of o-vanillin and RG-7112 reduced *p16^Ink4a^* expression in the IVDs of *sparc^-/-^* mice with established degeneration, supporting their ability to clear SnCs during late-stage disease (54). *P21* expression has not previously been examined in this context. We evaluated both *p16^Ink4a^*and *p21* to reflect different stages of the senescence; *p21* represents an early marker, while *p16^Ink4a^* represents a late and more stable marker of senescence (62, 63). As expected, *sparc*^-/-^ mice exhibited significantly higher levels of *p16^Ink4a^* and *p21*-positive cells in both NP and AF regions compared to *wt* mice (Figure 4E-K). Treatment with either o-vanillin or RG-7112 alone significantly reduced the number of *p16^Ink4a^* and *p21* positive cells in the AF and NP, with combination therapy resulting in the most pronounced reduction, approaching levels observed in *wt* controls (Figure 4 F-K).

**Figure 4.**
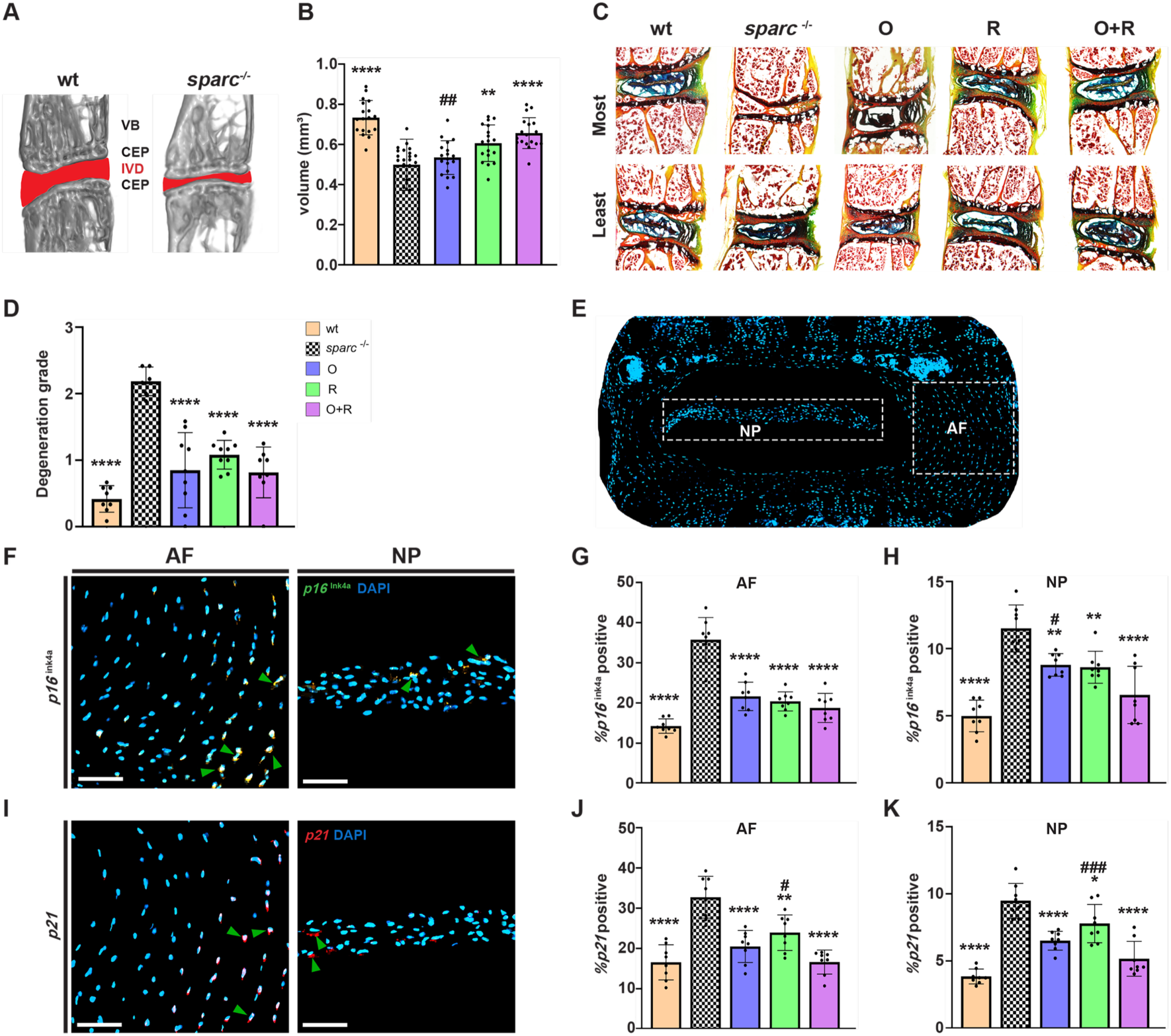
Senotherapeutic treatment increased disc volume, improved IVD integrity, and reduced senescent cell burden. **(A)** Representative micro-CT images outlining the IVDs (red) in 9-month-old *sparc*^-/-^ and wt mice. **(B)** Quantification of disc volume (L4–S1) (n = 6–7 animals per group; 3 males and 3–4 females). **(C)** Representative histological images of lumbar IVDs showing the least and most degenerated discs within each group. **(D)** Histological degeneration score assessed at three levels per animal (L4–S1), averaged, and reported for each group (n = 8–9 animals; 4–5 males and females). **(E)** Representative IVD section of NP and AF regions (dashed outlines). (**F)** Representative immunofluorescence staining of *p16^Ink4a^*, **(G-H)** Quantification of *p16^Ink4a^*-positive cells as a percentage of the total number, (**I)** Representative immunofluorescence staining of *p21*, **(J-K)** Quantification of *p21*-positive cells as a percentage of the total number in the AF and NP regions. DAPI served as a counterstain to measure the total number of cells. Statistical comparisons were calculated using an ordinary one-way ANOVA, with a Tukey’s post hoc analysis. Data are presented as means ± SD. */#*P* < 0.05, **/##*P* < 0.01, ###*P* < 0.001, \*\*\*\**P* < 0.0001. * Indicates a significant difference compared to *sparc*^-/-^, and # indicates a significant difference between single and combination treatment. Scales bars represent 100 um in F and I.

### Senotherapeutics improved health and reduced senescence in the cartilaginous endplates

Degeneration and senescence within the IVD have been extensively studied, whereas senescence and cartilage endplate (CEP) status have previously not been determined in the *sparc*^-/-^ model. The CEP is a critical structure regulating nutrient and metabolite transport between the disc and vertebral body (64, 65). Disruption of CEP integrity can impair disc homeostasis and exacerbate degeneration. We utilized micro-CT to investigate the effects of senolytic therapies on cartilage endplate health. In vehicle-treated *sparc*^-/-^ mice, bone volume fraction (BV/TV %) was significantly higher (Figure 5A-B), while endplate porosity was significantly lower in *sparc*^-/-^ compared to wt mice (Figure 5A, C). These changes may limit nutrient exchange and contribute to a hostile microenvironment for disc cells, thereby promoting degeneration and cellular senescence. Treatment with single drugs resulted in a lower BV/TV%, reaching statistical significance for RG-7112 compared to vehicle-treated *sparc*^-/-^ mice. The combination therapy resulted in the most pronounced and robust reduction in BV/TV%, significantly lower than single-drug treatments; however, BV/TV% values did not fully reach *wt* levels. Additionally, both single-drug treatments showed increased porosity, with the combination therapy resulting in the greatest increase in porosity, preserving structural integrity to levels comparable to *wt* mice (Figure 5C).

Next, to evaluate the effects of senolytic treatments on SnCs in the CEP, we quantified the number of *p16^Ink4a^* and *p21*-positive cells. Both markers were significantly elevated in *sparc*^-/-^ compared to wt mice (Figure 5I-J). Treatment with either o-vanillin or RG-7112 alone led to a lower number of *p16^Ink4a^*and *p21*-positive cells within the cartilage endplates. The expression of *p16*^Ink4a^ was not statistically different from wt in any of the treatment groups. Although *p21* was significantly lower in all groups, it only reached the level of *wt* following the combination treatment.

**Figure 5.**
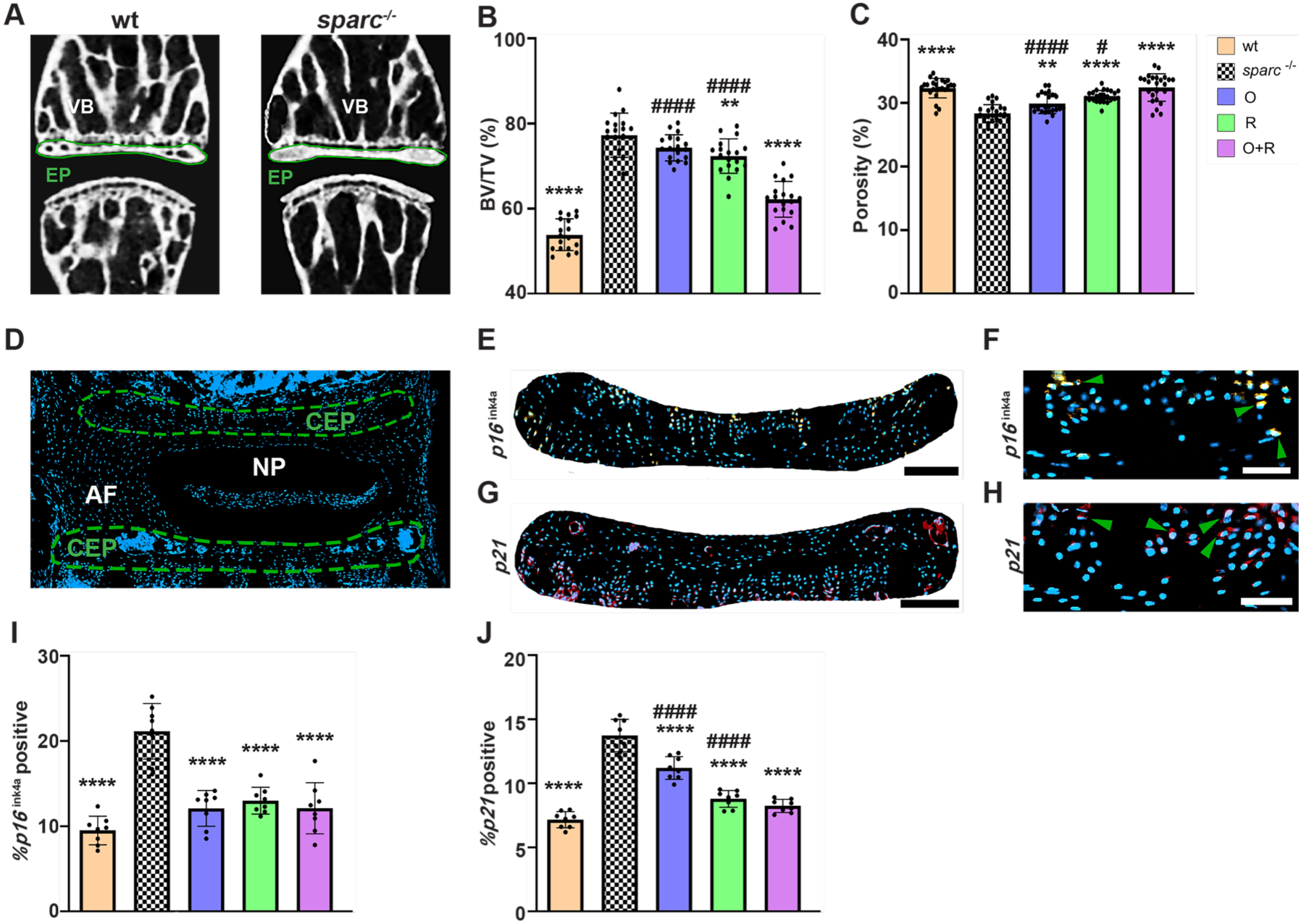
Senolytic treatment enhanced cartilage endplate structure and reduced senescent cell accumulation. **(A)** Representative micro-CT images with cartilage, the endplates outlined in green lines. **(B)** Bone volume over tissue volume (BV/TV %) and **(C)** porosity (%) was quantified in the CEPs (L4–S1) (n = 6 animals per group; 3 males and 3 females). **(D)** Representative IVD section illustrating NP, AF regions and the CEP region outlined by a green dashed line. Immunofluorescence staining of **(E-F)** *p16^Ink4a^* and **(G-H)** *p21* in the CEP. The green arrowheads indicate **(F)** *p16^Ink4a^* and **(H)** *p21* positive cells. Quantification of **(I)** *p16^Ink4a^* and **(J)** *p21* positive cells expressed as percentage of total cells. DAPI (blue) served as a nuclear counterstain. Data are presented as means ± SD and analyzed using one-way ANOVA followed by Tukey’s post hoc test. #*P* < 0.05, \*\**P* < 0.01, ****/####*P* < 0.0001. * Indicates a significant difference compared to *sparc*^-/-^, and # indicates a significant difference between single and combination treatment. Scale bars 100 um in E, F, G and H.

### Senotherapeutics improved bone parameters and reduced senescence in the vertebral bodies

In addition to IVD degeneration and LBP, *sparc*^-/-^ mice are known to have osteopenia (66, 67). As expected *sparc*^-/-^ mice displayed significantly lower bone quality compared to the *wt* group. The analysis revealed markedly reduced bone volume fraction (BV/TV %) (Figure 6B), trabecular thickness (Tb. Th) (Figure 6C), and trabecular number (Tb. N) (Figure 6D), accompanied by a significantly increased trabecular separation (Tb. Sp) (Figure 6E) in *sparc*^-/-^ mice compared to wt mice. Single-drug treatment with o-vanillin or RG-7112 significantly improved Tb. Th (Figure 6C). Combination treatment resulted in a robust and significant improvement in all parameters except Tb. Sp. Analysis of cortical bone parameters showed significantly lower bone volume (BV) (Figure 6F) and moment of inertia (MMI) (Figure 6 G) in *sparc*^-/-^ compared to wt mice. Single-drug and combination treatment significantly increased BV. Improvements in the MMI were observed following single-drug treatments, and the combination therapy did not provide an additive effect (Figure 6F-G). The Cs. Th parameter did not show a significant difference between the groups (Figure 6H). These findings demonstrate that oral administration of senolytic drugs, particularly in combination, prevents the early deterioration of bone quality in young *sparc*^-/-^ mice.

To evaluate the effects of senolytic treatment on senescence markers in the vertebral bone, we performed immunofluorescence staining for *p16*^Ink4a^ and *p21*-positive SnCs (Figure 6I-J). Both markers were significantly higher in the vertebral bone of *sparc*^-/-^ compared to *wt* mice. Treatment with single senolytic drugs led to a substantial reduction in the fluorescent intensity of both *p16^Ink4a^* and *p21 in* treated *sparc*^-/-^ mice. Combination therapy provided an even greater reduction in the markers compared to single drugs, suggesting a synergistic effect of the two drugs (Figure 6L-M).

**Figure 6.**
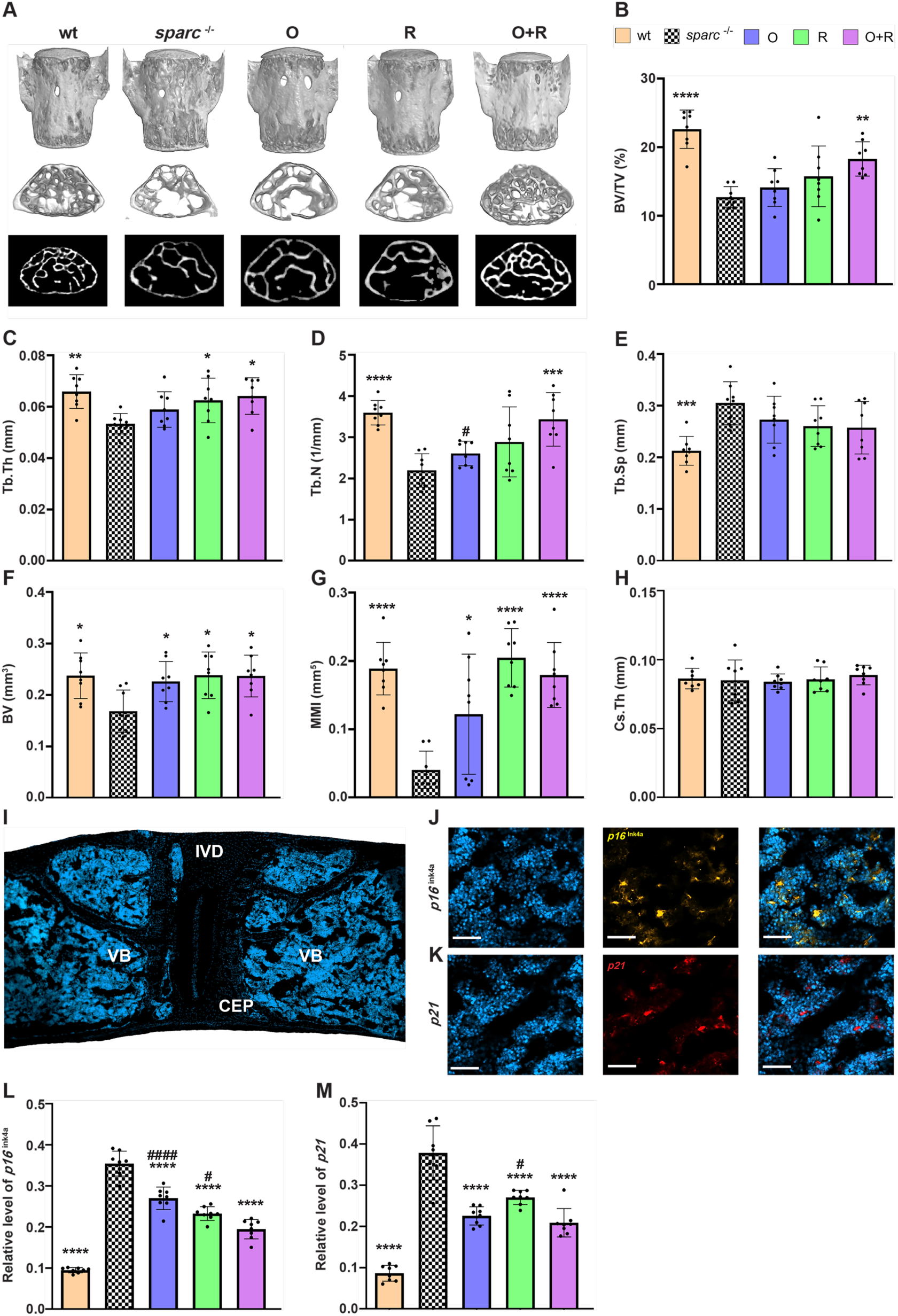
Senotherapeutics improved vertebral bone microarchitecture and decreased senescent cell burden. **(A)** Representative micro-CT images of trabecular and cortical bone in all groups. **(B–E)** Quantification of trabecular and **(F–H)** cortical bone parameters. **(I)** Representative vertebral bone section including IVD and CEP regions. **(J)** Immunofluorescence staining of *p16^Ink4a^* and **(K)** *p21* in vertebral bone. **(L, M)** Quantification of fluorescent intensity relative to the counterstain (DAPI). Data are presented as means ± SD and analyzed by one-way ANOVA with Dunnett’s post hoc test. */#*P* < 0.05, \*\**P* < 0.01, \*\*\**P* < 0.001, and \*\*\*\**P* < 0.0001. * Indicates a significant difference compared with *sparc*^-/-^group. N = 8 animals (4 males and 4 females) per group and 3 levels per animal (L4-S1). Scale bars 100 um in J and K. Quantification of *p16^Ink4a^*-positive cells in the dorsal horn. (**C**) Representative image showing*p16^Ink4a^*, and GFAP expression. **(D)** Percentage of GFAP and GFAP/ *p16^Ink4a^*double-labelled cells. **(E)** Representative image showing *p16^Ink4a^*and Iba1 expression. **(F)** Percentage of Iba1 and Iba1/ *p16^Ink4a^* double-labelled cells. **(G)** Representative image showing *p16^Ink4a^*, and NeuN expression. **(H)** Percentage of NeuN and NeuN/ *p16^Ink4a^* double-labelled cells. Number of cells/ mm^2^. N= 8-10 animals per group (4 to 5 males or females). DAPI served as a nuclear counterstain. For each animal, the mean of three independent images was calculated for group analysis. Scale bars = 200 μm and 80 μm. Data are presented as means ± SD and were analyzed by an ordinary one-way ANOVA followed by Tukey’s post hoc test. \**P* < 0.05, \*\**P* < 0.01, \*\*\**P* < 0.001, and \*\*\*\**P* < 0.0001.* Indicates a significant difference compared with *sparc^-/^*^-^.

## Discussion

Cellular senescence is increasingly recognized as a causal driver of musculoskeletal degeneration and chronic pain (3). Within the IVD, accumulation of SnCs promotes extracellular matrix breakdown, inflammatory signalling, and discogenic pain (42, 44, 68, 69). Senolytic strategies that modulate SASP production or selectively eliminate SnCs have shown therapeutic promise in age-related tissue dysfunction (70, 71). Because IVD degeneration is tightly linked to LBP, senotherapy offers a compelling opportunity for disease prevention (53, 54, 72, 73). Here, we demonstrate that systemic oral administration of o-vanillin and RG-7112 in young *sparc^-/-^* mice prevents behavioral manifestations of LBP, suppresses SASP activity, and reduces senescence markers across multiple spinal compartments. Treatment preserved disc volume, maintained vertebral and endplate microarchitecture, and reduced senescent astrocytes, microglia, and neurons in the dorsal horn, supporting the potential of senolytics to delay degenerative and pain-related cascades.

We previously established the senolytic properties of o-vanillin, a natural small molecule, and RG-7112, a synthetic MDM2 inhibitor (49–54, 74). Both agents selectively cleared SnCs and reduced SASP factors in senescent IVD cells, ex vivo human tissue, and middle-aged *sparc^-/-^* mice with advanced IVD degeneration (49, 53, 54). Given the heterogeneous phenotypes and survival mechanisms of SnCs, recent evidence suggests that combining senolytics with distinct molecular targets may enhance therapeutic outcomes (75–77). O-vanillin and RG-7112 engage different pathways associated with senescent cell survival. O-vanillin primarily modulates NF-κB and NRF-2–related signalling (49, 78), while RG-7112 activates p53-dependent apoptosis through MDM2 inhibition (51, 53). Our prior findings support that their combined use results in a synergistic effect, producing a more robust reduction in senescent cell burden and SASP expression across both in vitro and in vivo models of intervertebral disc degeneration when the degenerative milieu has already been established (51, 54).

*Sparc^-/-^* mice spontaneously develop IVD degeneration and pain by 4 months of age (58). Initiating senolytic treatment before the onset prevented the emergence of pain-like behaviours, with treated animals exhibiting sensitivity comparable to wt controls at 9 months. Both o-vanillin and RG-7112 provided measurable benefits, but combination treatment produced the most robust effects across behavioural assays. Prior studies have shown that SnC clearance mitigates cartilage and disc degeneration and reduces pain in osteoarthritis models (79–81). Our data extends these findings by demonstrating that early senotherapy interrupts the trajectory toward chronic pain, establishing senescence as a mechanistic link between degeneration and nociception.

Neuroinflammation is a critical component of chronic pain pathogenesis. We observed elevated *p16^Ink4a^* expression in the dorsal horn of *sparc^-/-^* mice, consistent with prior reports implicating spinal senescence in pain states (54, 61, 82, 83). SnCs accumulate in neurons, glia, and astrocytes in pain models and patient tissues (84–93), and their SASP promotes neuroinflammation and sensitization (61, 82, 88). Senolytic interventions in spinal cord injury reduce inflammation, glial scarring, and promote functional recovery (94). Consistently, we found senescent astrocytes, microglia, and neurons in the dorsal horn, all reduced by senotherapy. Unlike peripheral tissues, combination treatment did not show additive effects in the spinal cord, suggesting that CNS-resident SnCs may share common vulnerabilities. These findings highlight senescent neural and glial populations as critical contributors to central sensitization and support their removal as a strategy to restore spinal homeostasis and alleviate pain (61, 82, 88).

Senolytic treatment broadly suppressed SASP activity within IVDs. The IVDs of vehicle-treated *sparc^-/-^* animals displayed elevated levels of inflammatory cytokines and chemokines, whereas o-vanillin and RG-7112 treatment reduced all 15 factors measured. In contrast, interventions in middle-aged *sparc^-/-^*animals attenuated only 10 factors (54), suggesting that early therapy intercepts pro-inflammatory feedback before stabilization. Notably, chemokines such as CXCL-1, CXCL-5, and CXCL-10 linked to ECM degradation and senescence propagation (53, 95–97) were strongly reduced. Similarly, CCL-2 and CCL-7, mediators of immune recruitment and joint inflammation (52–54, 97), were suppressed. Key cytokines central to disc degeneration and osteoarthritis, including IL-1β, TNF-α, and IL-6, were also decreased, with IL-6 showing an additive reduction under combination treatment. Downregulation of VEGF-α and RANKL further indicates that early senotherapy limits angiogenic and osteoclastogenic cascades (98). Collectively, these results demonstrate that early intervention achieves more comprehensive inflammatory suppression than late treatment.

Histological analyses confirmed reductions in *p16^Ink4a^*-and *p21*-positive cells in IVDs, CEPs, and vertebral bone. Notably, clearance of *p21*-positive CEP cells required combination treatment, suggesting differential vulnerability among senescent subpopulations. This aligns with evidence that *p21*-driven senescence can resist single-agent therapy (99–101). Prior studies demonstrate enhanced efficacy of combination senolytics (45, 72, 102–108), and our findings reinforce this in disc and bone tissues. Importantly, our oral delivery route contrasts with previous intraperitoneal regimens (72), offering greater translational potential.

It is well established that the NP and the inner two-thirds of the AF lack direct vascular supply and instead rely on diffusion from capillaries located in the subchondral region of the endplate for nutrients (109, 110). With advancing age, senescence of vascular endothelial cells and osteoblasts within the endplate is thought to contribute to reduced bone mass and capillary density in this region, ultimately compromising nutrient delivery to the IVD (80, 111, 112). Targeting senescent cells in the bony endplate with senolytic therapy may help restore this nutrient pathway and support disc health. Prior studies in aged mice have reported reductions in IVD height associated with vertebral bone loss and marked disruption of endplate vasculature, changes that likely impair nutrient transport (111). Additionally, recent findings indicate that osteoclasts within the vertebral endplate undergo senescence in response to injury and aging, with elevated levels of TRAP, SA-βGal, and *p16^Ink4a^*observed in both aged and mechanically injured models (80). In our study, treatment with o-vanillin and RG-7112 significantly reduced the number of *p16^Ink4a^* and *p21*-positive cells in the cartilage endplate and increased its porosity. These changes suggest that senolytic intervention may improve structural properties of the endplate and facilitate nutrient diffusion into the IVD.

*Sparc^-/-^* mice display osteopenia, which is thought to contribute to the onset and progression of LBP (66, 113). Treated animals showed preserved vertebral trabecular and cortical architecture with reduced senescent osteocytes and osteoclasts, consistent with prior senolytic studies in skeletal aging (103, 106, 111, 114–116). Unlike ABT263, which may impair osteoblast differentiation (116), or D+Q, which showed variable effects in vertebrae (72), o-vanillin and RG-7112 preserved bone integrity, highlighting distinct therapeutic advantages.

Mechanistically, o-vanillin and RG-7112 target divergent survival pathways. RG-7112 induces apoptosis in p53/p21-driven SnCs, whereas o-vanillin modulates NF-κB and *p16^Ink4a^*-driven populations (49, 53, 54). Additive suppression of IL-6 and enhanced clearance of *p21*-positive cells support complementary activity. Importantly, treatment timing strongly influenced outcomes. In middle-aged *sparc^-/-^*mice, both agents were required at full dose, and only combination treatment achieved comprehensive effects (54). In contrast, early intervention allowed effective suppression with monotherapy, indicating greater vulnerability of nascent SnCs.

In summary, our data establish that oral senotherapy with o-vanillin and RG-7112 prevents senescent cell accumulation, suppresses SASP production, preserves disc and bone integrity, and prevents chronic pain behaviours in *sparc^-/-^*mice. These findings identify cellular senescence as a mechanistic driver of spine degeneration and pain and demonstrate that early intervention provides durable protection across multiple spinal compartments. The results support translation of senotherapeutic strategies toward preventive clinical applications in individuals at risk for early-onset or genetically driven disc degeneration.

## MATERIALS AND METHODS

### Study design

This study aimed to evaluate the preventive effects of the senolytic compounds o-vanillin and RG-7112 on IVD degeneration and associated LBP. We employed the *sparc*^-/-^ mouse model, which we have previously demonstrated to exhibit early-onset of IVD degeneration, back pain and increased accumulation of SnCs in the spine (54). Based on this phenotype, we assessed whether early senolytic treatment could prevent or reduce the accumulation of SnCs and preserve spine health. To further characterize the effect of senolytic treatment on pain modulation, pain behaviour was assessed monthly using grip strength, von Frey, and acetone-evoked tests. We also performed immunostaining for markers of activated astrocytes, microglia, and neurons across spinal tissues from untreated wt mice and both treated and untreated *sparc*^-/-^ mice. At the endpoint, lumbar IVDs were harvested for SASP factor analysis by Luminex multiplex assay. SnC depletion was evaluated by *p16^Ink4a^* and *p21* immunofluorescence in IVDs, cartilage endplates, vertebral bone, and spinal cord. Structural improvements in discs and bone were assessed by micro-CT for disc volume, endplate porosity, and bone microarchitecture, and by FAST histological grading for IVD degeneration.

### Animals and housing

All animal procedures were approved by the McGill University Animal Care Committee in accordance with the Canadian Council on Animal Care guidelines (Protocol number: MUHC-10007). Age-matched male and female C57BL/6N wild-type and *sparc*^-/-^ mice were used across all experimental groups. The *sparc*^-/-^ mouse model, originally generated on a C57BL/6×129SVJ background, was maintained through successive backcrossing to the C57BL/6N strain (59, 117).

Mice were housed in groups of two to four per cage in ventilated polycarbonate cages (Allentown) under standard conditions (12-hour light/dark cycle, controlled temperature and humidity). Environmental enrichment was provided via cotton nesting material, and cages were lined with corncob bedding (Envigo). Animals had ad libitum access to irradiated, soy-free extruded rodent chow and filtered water.

Group sizes were determined based on prior data and power calculations from previous experiments involving *sparc*^-/-^ mice, ensuring sufficient statistical sensitivity to detect both genotype and treatment-related effects (54, 118–123). For cytokine profiling using the Luminex multiplex assay, six animals per group were utilized. For all other experimental outcomes, including behavioral, imaging, histological, and molecular analyses, group sizes ranged from eight to fifteen animals per condition.

### Randomization and blinding

Animals were randomly assigned to each treatment group. Experimenters were blinded to genotype and treatment for all experiments and data analysis.

### Treatment regime and endpoints

Four-month-old male and female mice were randomly assigned into five treatment groups. Both *sparc*^-/-^ and wt mice received oral gavage weekly for a duration of 5 months. *Sparc*^-/-^ mice were treated with one of the following senolytics: o-vanillin (100 mg/kg), RG-7112 (5 mg/kg), a combination of both drugs (o-vanillin + RG-7112), or vehicle control (0.01% dimethyl sulfoxide in saline). Wild-type mice received only vehicle treatment to serve as baseline controls.

The frequency and dosage of drug treatment, as well as the experimental endpoints, were determined based on previous studies (54, 118–123). The initiation of treatment was chosen based on the onset of the pain phenotype and IVD degeneration. *Sparc*^-/-^ mice begin to exhibit signs of back pain and disc degeneration at 4 months of age (57–59). Our objective was to assess whether senolytic treatment, with o-vanillin and RG-7112, administered either as a single or in combination, could provide a protective effect against the progression of these phenotypes.

### Pain behavior

Pain-related behaviour was evaluated as was previously described (54, 58, 59, 119, 124, 125). All behavioural testing was conducted in a dedicated room under standard indoor lighting between 8:00 a.m. and 12:00 p.m. To reduce stress, mice were habituated to the testing environment for 1 hour, followed by an additional 1-hour habituation period in Plexiglas testing boxes placed on a metal mesh platform when applicable. Each behavioural test lasts between 30 s and 5 min per animal. The animals were subjected to one test per day, except for the first baseline measurements, when von Frey and acetone were tested on the same day. Animals were closely monitored for signs of distress or discomfort during testing before being returned to the animal facility.

#### Grip strength

Axial discomfort was assessed using a grip strength meter (Stoelting Co.). Mice were encouraged to grasp the bar with their forepaws while gentle, steady traction was applied via the tail until they released their grip. The maximal force was recorded in grams (123). For each session, grip strength was measured three times per mouse, and the values were averaged to obtain a final score. To avoid fatigue and stress, animals were returned to their home cages for approximately 15 minutes between measurements.

#### Acetone-evoked behavior test for cold sensitivity

Behavioral reaction to a cold stimulus was used to determine radiating pain. Acetone (30 to 50 μl) was applied to the left and right hind paws, and the total duration of nocifensive behavior (paw lifting, shaking, and scratching) was recorded for 30 s (123).

#### Von Frey test for mechanical sensitivity

Mechanical sensitivity was assessed using von Frey filaments (Stoelting Co.) applied to the plantar surface of the left and right hind paws. Each filament was applied with enough force to slightly bend the filament and held in place until a withdrawal response occurred or for a maximum of 5 seconds, whichever came first (54, 59). Stimulus intensity ranged from 0.6 to 4.0 g, corresponding to filament numbers 3.22, 3.61, 3.84, 4.08, and 4.17. The 50% paw withdrawal threshold was calculated using the up-down method (126), providing a quantitative measure of mechanical hypersensitivity.

### Luminex multiplex assay

SASP factor release from the IVDs was evaluated at the termination of treatment. Animals were deeply anesthetized via intraperitoneal injection of a mixture containing ketamine (100 mg/kg), xylazine (10 mg/kg), and acepromazine (3 mg/kg). Transcardial perfusion was performed using a vascular rinse, and tissues fixation by 4% paraformaldehyde (PFA) in 0.1 M phosphate buffer (PBS) (pH 7.4) at room temperature. Lumbar IVDs (L1–L5) with intact cartilaginous endplates, without vertebral bone were dissected for analysis. The IVDs were maintained for 48 hours in DMEM supplemented with 1×GlutaMAX, penicillin (10 U/mL), and streptomycin (10 μg/mL) in an incubator (54). The conditioned medium was then collected for protein analysis. The concentration of 15 proteins (CXCL-1, CXCL-5, CXCL-9, CXCL-10, CCL-2, CCL-7, IL-1β, IL-2, IL-6, IL-10, TNF-α, IFN-γ, VEGF-α, RANKL, and M-CSF) were measured using a Luminex multiplex assay (Thermo Fisher Scientific, PPX-15-MXGZF4V), according to the manufacturer’s instructions. Data acquisition was performed using the MAGPIX system with xPONENT software, and results were processed and analyzed using the ProcartaPlex Analysis App.

### Histological analysis

#### Sample preparation

Animals were deeply anesthetized and transcardial perfusion was performed as described under the “Luminex multiplex assay” section. Following perfusion, the T13–S1 spinal segment was harvested and post-fixed overnight in 4% PFA at 4°C. Samples were then decalcified in 4% ethylenediaminetetraacetic acid (EDTA) in PBS at 4°C for 14 days. After decalcification, tissues were cryoprotected in 30% sucrose in PBS for 4 days at 4°C, then embedded in optimal cutting temperature (OCT) compound (Tissue-Tek, Sakura Finetek, Torrance, CA, USA). Sagittal sections were cut at 16 μm thickness using a cryostat (Leica CM3050S, Leica Microsystems Inc., Concord, Ontario, Canada), thaw-mounted onto gelatin-coated glass slides, and stored at −20°C until further processing.

#### FAST staining

Staining was performed on sagittal spinal sections (three per animal) using the FAST protocol, as previously described by Millecamps et al. (54, 57–59, 120, 124, 125). IVD degeneration severity grading scale: Grade 0: Healthy IVDs display intact structure, a clear distinction between outer AF and inner NP and negatively charged proteoglycans; grade 1: The changes in extracellular components and IVD integrity were identified as grade 0, normal structure, but the loss of proteoglycans in inner NP; grade 2: internal disruption (loss of boundary) between NP and AF; grade 3: bulging of NP in dorsal aspect; and grade 4: herniation. Each value represents the average grading score of 3 to 4 IVDs per animal.

### Spine, endplate, bone and spinal cord immunofluorescent histochemistry

Sagittal spinal sections (three per animal) were processed for immunofluorescent detection of senescence markers following the protocol previously described by Mannarino et al. (54). Additionally, three spinal cord sections per animal were randomly selected, spanning the lumbar spinal cord for each antibody. Sections were incubated in blocking buffer for 1 hour at room temperature. Slides were then incubated with appropriate antibodies (Table S2) in blocking buffer overnight at 4°C. After PBS washes, sections were incubated for 1.5 hours at room temperature with appropriate secondary antibodies (Table S2) in blocking buffer. DAPI (1:50,000 in PBS; Sigma-Aldrich) was briefly applied, and slides were washed another three times for 5 min. Coverslips were mounted using Aqua-Poly/Mount (Polysciences Inc.). Fluorescence images were captured at 10× magnification using an Olympus BX51 microscope equipped with an Olympus DP71 camera. For the spinal cord, colocalization of *p16^Ink4a^* with GFAP, Iba1, and NeuN in the dorsal horn was quantified using ImageJ. The number of *p16^Ink4a^*-positive cells was expressed as the number of double-labelled cells per mm² relative to each marker (GFAP, Iba1 and NeuN). For the IVD and endplate regions, the percentage of *p16^Ink4a^* or *p21*-positive cells was calculated by dividing the number of positively stained nuclei by the total number of DAPI-stained nuclei. Approximately 5,000-7000 cells were quantified per treatment group. For vertebral bone sections, *p16^Ink4a^* or *p21* signal intensity was quantified within a defined region of interest (ROI; 1000 μm²) located immediately below the growth plate. A threshold was set to distinguish specific immunoreactivity from background, and the relative signal was calculated by normalizing *p16^Ink4a^*or *p21* immunoreactivity to DAPI. The average percent area of immunoreactivity across the three sections per animal was used as a single biological replicate. All image quantification was performed by three independent experimenters blinded to both genotype and treatment group.

### Micro-Computed Tomography (Micro-CT)

Micro-CT imaging and analysis were performed as previously described (54, 111, 127). High-resolution scans of fixed spines were acquired from the L4 to S1 vertebral levels to evaluate three-dimensional (3D) structural features. Scans were conducted using a Skyscan 1172 micro-CT system (Bruker, Kontich, Belgium) with a spatial resolution of 15 μm. The system was equipped with a 0.5-mm aluminum filter and operated at 45 kV and 220 μA with a 360° rotation. Images were captured at 0.4° rotation steps, averaging four frames per image, with an exposure time of 1.46 seconds per frame. Image reconstruction was performed using the nRecon software (v1.7.1.0, Bruker), and quantitative analysis was carried out using CTAn software (v1.18.8.0, Bruker). CTVOX (v3.3, Bruker) was used for 3D visualization of bone morphology and disc architecture. Transverse cross-sectional images were analyzed to assess disc volume and bone structure. Disc volume was measured by manually defining a ROI encompassing the space between adjacent endplates. For trabecular bone analysis, the ROI was delineated between the endplate and transverse process within the vertebral body. From these 3D datasets, standard trabecular bone parameters were calculated, including bone volume fraction (BV/TV), trabecular thickness (Tb. Th), trabecular number (Tb. N), and trabecular separation (Tb. Sp). Cortical bone parameters were assessed using two-dimensional (2D) measurements, including cortical bone volume (BV), cortical thickness (Cs. Th), and polar moment of inertia (MMI).

### Statistical analysis

Power analysis was conducted with an alpha (α) level of 0.05, a confidence level of 95%, and a 50% response distribution. Based on these parameters, a sample size of 6 for Luminex assay and 8 to 10 animals for the other experiments per group (wt and *sparc^-/-^*) was determined to be sufficient to detect statistically significant differences, with 10 to 15 animals required if sex-specific effects were present. All statistical analyses were performed using GraphPad Prism 10. Data are presented as mean ± standard deviation (SD), and significance was set at P ≤ 0.05. Depending on the experimental design, comparisons were made using two-tailed unpaired t tests, repeated-measures one-way ANOVA, or repeated-measures two-way ANOVA, followed by Dunnett’s or Tukey’s post hoc tests where appropriate.

