## Supplementary Figures and Tables for "Senolytic Therapy as a Preventive Strategy for Low Back Pain"

Saber Ghazizadeh *et al.*

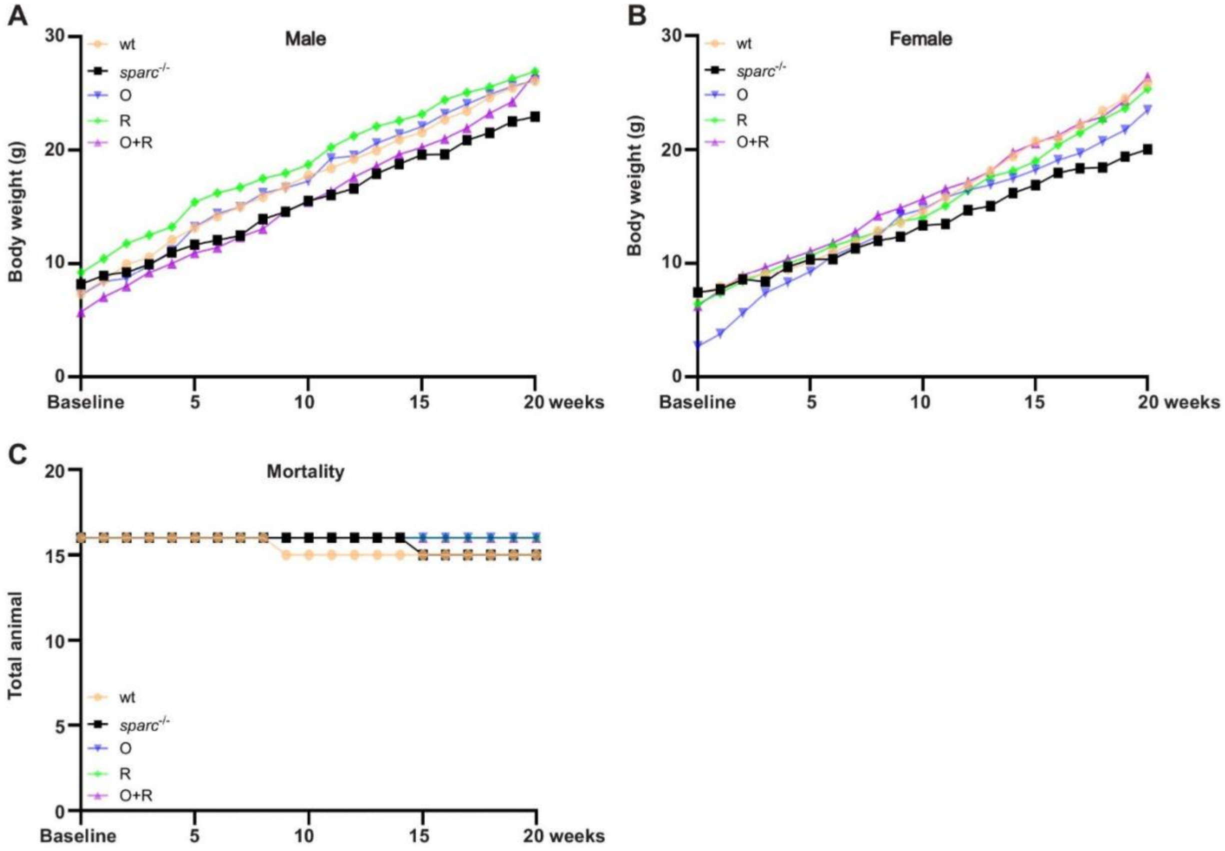


**Figure S1. Senolytic treatment does not affect body weight and mortality.**

**(A)** Body weight of male and **(B)** Female mice in all groups. **(C)** Mortality rates at baseline up to 20 weeks of wildtype, *sparc*$^{-/-}$ and treatment groups. Data was presented as mean ± SD and analyzed by one-way ANOVA followed by Tukey's post-hoc test. n = 7- 10 animals (4-5 male and 3-5 female) per group, treated with senolytics or vehicles.

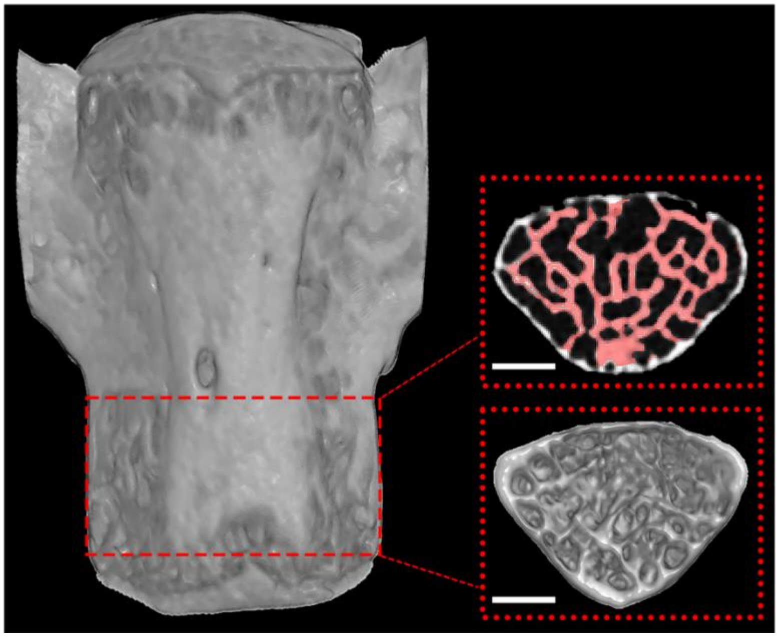

**Figure S2. Spine reconstruction.** Representative image of the ROI (red dashed line) selected to measure cortical and trabecular bone parameters. The bottom area indicates the endplate, and the upper area indicates the transverse processes, which were landmarks for the ROI selection.

**Table S1.** SASP average concentrations (pg/mL) following ex vivo treatment with Senotherapeutics (Average ± SD)

| | **wt** | ***sparc$^{-/-}$*** | **O** | **R** | **O+R** |
|---|---|---|---|---|---|
| **CXCL1** | 123.28 ± 43.72 | 457.07 ± 118.5 | 221.22 ± 33.0 | 265.33 ± 77.14 | 218.74 ± 51.34 |
| **CXCL5** | 156.94 ± 37.63 | 654.57 ± 113.24 | 308.59 ± 63.18 | 366.43 ± 57.44 | 319.66 ± 50.27 |
| **CXCL9** | 370.88 ± 110.02 | 1700.49 ± 397.56 | 746.91 ± 378.65 | 629.18 ± 129.07 | 405.27 ± 76.62 |
| **CXCL10** | 47.32 ± 23.07 | 285.78 ± 77.79 | 82.13 ± 49.12 | 65.38 ± 10.47 | 40.15 ± 15.81 |
| **CCL2** | 2474.48 ± 769.29 | 8304.62 ± 1174.64 | 4472.66 ± 1336.84 | 5479.90 ± 904.40 | 4908.99 ± 602.63 |
| **CCL7** | 409.34 ± 119.79 | 1217.89 ± 101.03 | 568.89 ± 112.13 | 629.48 ± 20.06 | 591.48 ± 32.59 |
| **IL-1β** | 15.08 ± 4.19 | 73.38 ± 14.72 | 30.44 ± 7.93 | 34.46 ± 4.29 | 26.59 ± 3.87 |
| **IL-2** | 63.07 ± 11.49 | 206.22 ± 31.79 | 93.36 ± 15.21 | 105.33 ± 3.83 | 90.57 ± 5.89 |
| **IL-6** | 6160.36 ± 2510.03 | 58669.01 ± 4279.53 | 37887.63 ± 5028.09 | 41284.99 ± 4656.25 | 31952.18 ± 4740.37 |
| **IL-10** | 85.17 ± 24.17 | 374.36 ± 58.83 | 164.27 ± 26.87 | 187.50 ± 16.17 | 156.90 ± 16.17 |
| **TNF-α** | 139.54 ± 31.32 | 530.48 ± 68.38 | 230.07 ± 31.38 | 267.72 ± 29.44 | 226.68 ± 22.60 |
| **IFN-γ** | 12.30 ± 3.24 | 51.51 ± 9.73 | 25.19 ± 6.78 | 29.30 ± 3.06 | 22.77 ± 2.51 |
| **VEGF-α** | 231.96 ± 57.20 | 879.69 ± 148.04 | 314.14 ± 51.76 | 297.41 ± 66.79 | 220.01 ± 34.75 |
| **RANKL** | 18.04 ± 5.28 | 66.79 ± 13.33 | 29.71 ± 5.58 | 30.33 ± 3.36 | 25.54 ± 2.06 |
| **M-CSF** | 4.71 ± 0.79 | 15.63 ± 2.77 | 6.28 ± 1.49 | 5.59 ± 0.66 | 5.00 ± 0.65 |

**Table S2.** List of antibodies used in the study.

| Antibody | Source | Catalog Number | Dilution |
|---|---|---|---|
| Anti-CDKN2A/p16$^{INK4a}$ | Rabbit | ab211542 | 1:100 |
| Anti-p21 | Rabbit | ab188224 | 1:500 |
| IBA1 Monoclonal | Mouse | MA5-27726 | 1:100 |
| Anti-GFAP | goat | SAB2500462 | 1:500 |
| Anti-NeuN purified | guinea pig | ABN90P | 1:500 |
| Alexa Fluor™ 594 Donkey anti-Goat IgG | Goat | A-11058 | 5 µg/mL |
| Alexa Fluor® 488 Donkey Anti-Rabbit IgG (H+L) | Donkey | 711-545-152 | 1:2000 |
| Cy™3 Donkey Anti-Rabbit IgG (H+L) | Donkey | 711-165-152 | 1:2000 |
| Goat anti-Rabbit IgG (H+L), Cyanine3 | Goat | A10520 | 5 µg/mL |
| Alexa Fluor™ 647 Donkey anti-Rabbit IgG | Donkey | A-31573 | 5 µg/mL |
